# SLC25A39 loss disrupts mitochondrial redox homeostasis and exposes therapeutic vulnerabilities in glioblastoma

**DOI:** 10.64898/2026.11.12.765851

**Authors:** Katherine L Eales, Yingying Lyu, Yuan Feng, Ishana Rajgopal, Suvagata Roy Chowdhury, Bryan Marzullo, Harriet Palmer-Coulson, Colin Nixon, Jennie Roberts, Victoria Wykes, Julien Prudent, Wei Hua, Daniel A Tennant

**Affiliations:** Department of Metabolism and Systems Science, School of Medical Sciences, College of Medicine and Health, University of Birmingham, Edgbaston, Birmingham. B15 2TT. UK; Medical Research Council Mitochondrial Biology Unit, University of Cambridge, Cambridge, UK; Cancer Research UK Scotland Institute, Switchback Road, Bearsden, Glasgow. UK; Department of Cancer and Genomic Science, School of Medical Sciences, College of Medicine and Health, University of Birmingham, Edgbaston, Birmingham. B15 2TT. UK; Department of Neurosurgery, Queen Elizabeth Hospital Birmingham, University Hospitals Birmingham NHS Foundation Trust, Birmingham, B15 2GW. UK; Université Paris Cité, Institut Imagine, Laboratory of Mitochondrial Dynamics, INSERM UMR 1163, Paris, France; Department of Neurosurgery, Zhongshan Hospital, Shanghai Medical College, Fudan University, PR China; Sycamore Research Institute of Life Sciences, PR China

## Abstract

Glioblastoma is an incurable brain tumour characterised by extensive hypoxia and phenotypic plasticity, both of which drive therapy resistance. Mitochondrial glutathione, dependent on the inner membrane carrier SLC25A39, is central to buffering the oxidative stress that hypoxia generates; its regulation and significance in glioblastoma are nonetheless unknown. We find that SLC25A39 expression is hypoxia-inducible and HIF-dependent, enriched in hypoxic tumour niches and heterogeneously expressed within and between tumours. Its loss depletes mitochondrial glutathione, causing oxidative damage to nucleic acids and lipids, mitochondrial hyperfusion, impaired respiration and a compensatory shift towards glycolysis, glutamine catabolism and serine synthesis. SLC25A39-deficient cells grow poorly, yet resist temozolomide while becoming markedly sensitive to navitoclax, an inhibitor of Bcl-2 and Bcl-xL, which increases cell killing when combined with radiation and temozolomide. In this work, we identify SLC25A39 as a metabolic vulnerability and candidate stratification marker in glioblastoma.

## Introduction

Glioblastoma (GBM) is characterised by significant phenotypic plasticity, causing therapy resistance and poor patient outcomes. This plasticity is now understood to be driven by microenvironmental conditions in the tumour, such as low oxygen (hypoxia) and nutrient stress^1, 2^.

Much like many solid tumours, GBM have well-described significant hypoxic regions^3^, with some parts even tending towards anoxia^4, 5^ resulting in characteristic intratumoural necrosis. Hypoxia is a pro-malignant cell stress, driving phenotypic change through multiple mechanisms. Many of these are orchestrated by Hypoxic Inducible Factor (HIF) 1 and 2, heterodimeric transcription factors formed of a hypoxia-stabilised alpha subunit and a constitutively expressed beta subunit. Upon binding hypoxia-response elements (HREs), they have been shown to regulate the transcription of genes includes in tumour cell differentiation, angiogenesis, immune recognition and metabolism, among others^6, 7^. Importantly, hypoxia is one of the key drivers of cancer therapy resistance, occurring through multiple mechanisms including evasion of anti-tumour immunity, increased resistance to apoptosis, upregulation of multi-drug efflux pumps and reduction of radiation-induced radical formation on DNA^8, 9^.

Hypoxia also drives cancer cells to alter their metabolism through both transcriptional changes and direct inhibition of oxygen-dependent processes, such as respiration. Indeed, upon decreased NADH oxidation downstream of oxygen restriction, cells must increase the generation of ATP through glycolysis to compensate^10^. However, pyruvate oxidation to fuel the TCA cycle is further inhibited by the HIF1-mediated stabilisation of PDK1^11, 12^. Indeed, many of the transcriptional changes that alter mitochondrial activity in hypoxia have been suggested to represent an attempt to limit the production of reactive oxygen species (ROS) under these conditions. Despite this, hypoxia drives increased mitochondrial ROS generation^13^, with the potential to increase the rate of nuclear DNA damage without corresponding changes in the cellular antioxidant capacity^14^.

Cells use both enzymatic and non-enzymatic antioxidant systems to detoxify mitochondrial ROS before they reach the nucleus. Many of the enzymes that reduce cellular peroxide rely on NADPH or reduced glutathione for their activity^15^. Glutathione is synthesised exclusively in the cytoplasm, and therefore requires specific transporters for it to reach the mitochondrial matrix and exert its antioxidant functions in this key compartment. Two members of the SLC25 transporter family were recently identified as being required for this transport; SLC25A39 and SLC25A40^16, 17^. Interestingly, while both appear to be responsible for mitochondrial accumulation of glutathione in different cell types^18^, only SLC25A39 contains the matrix cysteine residues that enable iron-sulfur cluster-dependent stabilisation and protection from constitutive degradation by the protease AFG3L2^19, 20^. This post-translational regulation establishes SLC25A39 as the predominant mitochondrial transporter, dynamically responsive to both GSH availability and iron status^19, 20^. Interestingly, AFG3L2 activity itself has been shown as upregulated under hypoxia^21^, thereby providing a route through which hypoxia may alter the ability of mitochondria to transport glutathione.

Here, we identify SLC25A39 as being upregulated in hypoxia, and importantly in the hypoxic niche observed in GBM. We demonstrate that SLC25A39 is vital for maintaining mitochondrial redox homeostasis and that loss of SLC25A39 leads to significant mitochondrial oxidative damage and broader metabolic perturbations. Furthermore, we suggest a role for SLC25A39 in response of glioblastoma to treatment: while its loss impairs growth, it also induces a metabolic phenotype that promotes resistance to the key chemotherapy used to treat GBM, temozolomide (TMZ). However, the same cells show sensitivity to agents that target mitochondrial stress downstream of SLC25A39 deficiency. We therefore suggest that SLC25A39 is a novel metabolic vulnerability in GBM, and a potential route to allow for patient treatment stratification.

## Results

### SLC25A39 expression is heterogenous within and between glioblastoma and upregulated in hypoxic niches

Previous studies have shown that SLC25A39 expression is regulated by both mitochondrial GSH availability and AFG3L2 protease activity^19, 20^, the latter of which is hypoxia regulated. We therefore examined whether in GBM cells, hypoxia altered the expression of this inducible mitochondrial transporter. Surprisingly, we found that GBM-relevant hypoxia^5^ (0.3% oxygen) for 24 hours led to differential stabilisation of SLC25A39 in GBM cell models (Fig.1A). We hypothesised that, similarly to other hypoxia responses, the hypoxia-induced change in SLC25A39 expression may vary between cell models. We therefore tested 8, 24 and 48 hours of incubation in hypoxia, and found that indeed this was the case, with all lines tested, including primary GBM models, demonstrating SLC25A39 stabilisation in hypoxia (Fig. 1B and S1A). Given this unexpected result, we examined the upstream regulatory region of SLC25A39 for hypoxia response elements (HREs), the means by which the HIF heterodimer binds and induces transcription of its target genes. We confirmed the presence of multiple canonical HREs in the genomic region surrounding the transcription start site (Fig. S1B), suggesting that hypoxia-induction of transcription is indeed HIF-mediated. We therefore knocked down HIF-1α and HIF-2α using RNAi, and showed reduction of SLC25A39 stabilisation in the absence of either alpha subunits (Fig.1C and D).

**Figure 1:**
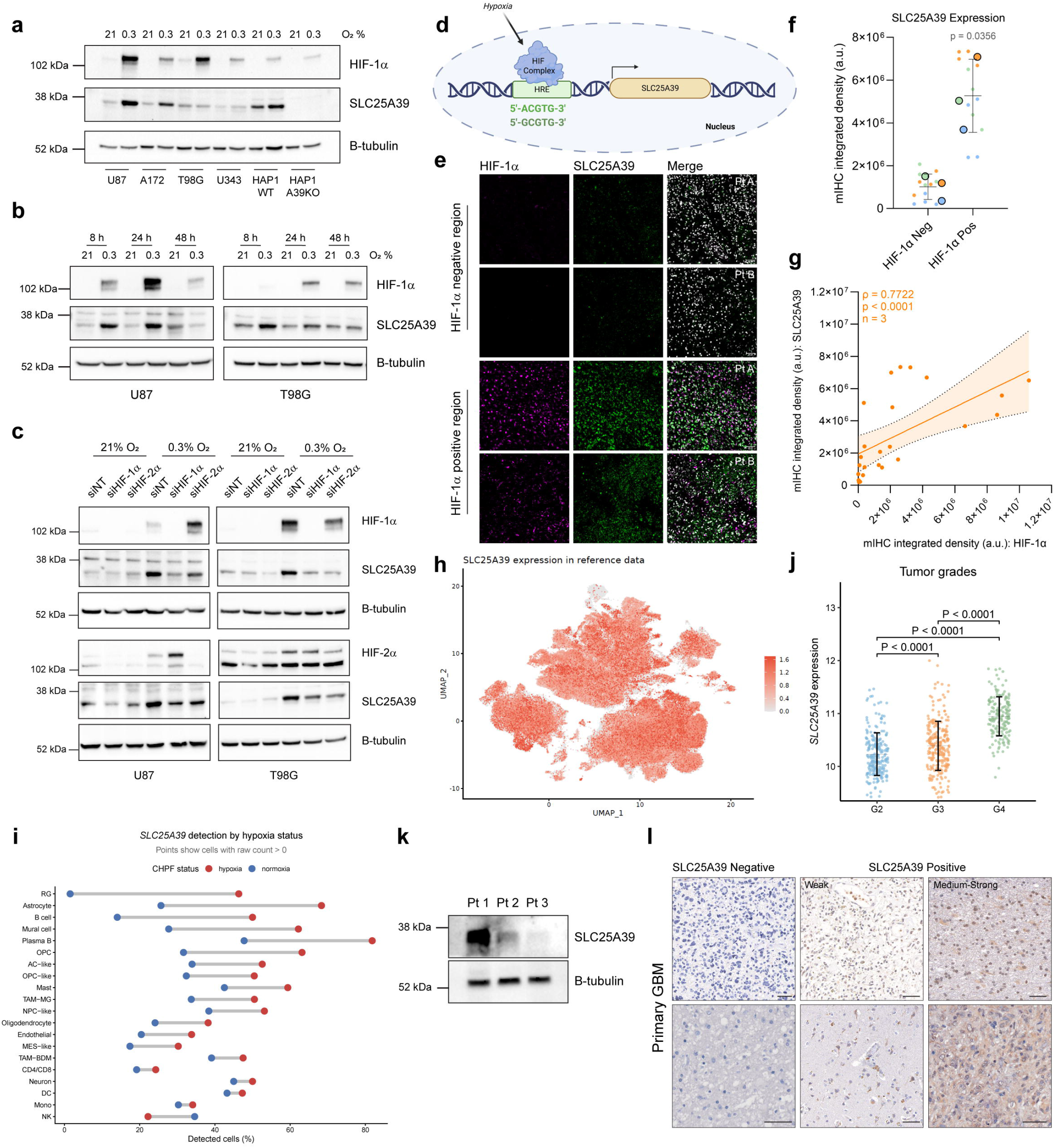
SLC25A39 expression is upregulated under hypoxic conditions. a) Representative western blot of SLC25A39 expression in glioma (U87, A172, T98G and U343) and HAP1 SLC25A39^+/+^ and SLC25A39^-/-^ cell lines under normoxia (21% O_2_) or hypoxia (0.3% O_2_) for 24 hours. b) Representative western blot of SLC25A39 expression in U87 and T98G cells at 21% or 0.3% O_2_ for 8, 24 and 48 hours. c) Representative western blot of SLC25A39 expression in control (siNT), siHIF-1α or siHIF2α U87 and T98G cells incubated in hypoxia for 8 hours (T98G) or 24 hours (U87). d) Schematic of hypoxia-responsive element (HRE) motifs in the upstream region of *SLC25A39*. Created in BioRender (https://BioRender.com/jbzbrh9). e) GBM tissue sections from two patients (Pt A, Pt B) showing SLC25A39 expression in areas with low and high HIF-1α expression. Scale bar = 50 µm. f) Multiplex immunohistochemistry (mIHC) quantified by integrated density in HIF-1α negative (Neg) and positive (Pos) areas of GBM tissue. Large data points are means from 3 patients; small points are individual regions of interest (ROIs; 4 per region, *n*=3 patients). g) Spearman’s correlation (ρ) between SLC25A39 and HIF-1α expression per ROI (n=3 patients). Orange line, linear regression; shaded band, 95% confidence interval. h) UMAP of SLC25A39 expression across all cells in GBMAP dataset. Colour intensity represents log-normalised expression. i) SLC25A39 detection rates in Cellular Hypoxia Prediction Framework (CHPF)-classified hypoxic versus normoxia cells by cell type. Detection defined as raw UMI count > 0. Blue and red points indicate normoxia and hypoxic percentages respectively. j) *SLC25A39* expression across glioma grades (grade 2-4) in the TCGA-LGG & GBM cohort. *P*-values are from pairwise Wilcoxon rank-sum tests. k) Western blot of SLC25A39 expression in cells from three patient tumours (Pt 1, Pt 2 Pt 3). l) Representative immunohistochemistry of SLC25A39 expression in primary GBM tissue. Scale bar = 50 µm. Western blots are representative of (**a**) *n* > 3 per cell line; (**b**) *n* = 4; (**c**) *n* = 4 (U87) and *n* = 3 (T98G) independent experiments. Data are means; error bars, SD.

Given that SLC25A39 stabilisation can occur in hypoxia in GBM cells, we investigated whether HIF-1α and SLC25A39 expression are correlated in GBM tumours. Comparing multiple tumour regions SLC25A39 showed significant variation across the cell mass, but was observed to be significantly higher in hypoxic, HIF-1α -positive areas (Fig. 1E-G and S1C), consistent with the in vitro data. We further analysed publicly available datasets to examine how expression of SLC25A39 mRNA varied between cell types within GBM. UMAP projections of cell type using the GBMap reference dataset suggested that it was detectable across all cells, but with significant variation (Fig. 1H, S1D and S1E). CHPF-predicted hypoxia status of GBM cell types showed that mesenchymal-like (MES-like) were most frequently hypoxic (Fig. S1F-H) and that hypoxia increased *SLC25A39* mRNA in all cell types, with the exception of the small NK cell population (Fig. 1I). We further extended this analysis to investigate whether bulk tumour mRNA levels of *SLC25A39* correlated with other aspects of tumour phenotype in the GDC/TCGA-GBM cohort. We first noted that *SLC25A39* mRNA increased with tumour grade (Fig. 1J). Interestingly, further analysis suggested that the correlation between *SLC25A39* and well-described hypoxia-responsive genes is increased with tumour recurrence (Fig. S1I and J). Notably, increased correlations were observed in for *P4HA1* and *BNIP3L*, genes involved in hypoxic adaptation and cellular stress, as well as *ANXA2* and *SDC2*, which have established roles in GBM invasion and extracellular matrix remodelling (Fig. S1I). These suggest that *SLC25A39*-associated transcriptional networks in recurrent GBMs are characterised by hypoxia-responsive, pro-invasive pathways.

Given previous classification of GBM subtypes^22^, we also investigated whether *SLC25A39* mRNA varied in this respect, observing a small but significant decrease in mesenchymal tumours, although this trend was reversed in recurrent GBM subtypes (Fig. S2A). Since *SLC25A39* mRNA expression increased with tumour grade, we evaluated whether expression correlated with patient overall survival. We found that there was no significant difference in all GBM, with a survival effect only detected when tumours were broken down into subtypes (Fig. S2B-D). Finally, as SLC25A39 protein may not correlate with mRNA due to the activity of AFG3L2, we extended the analysis of SLC25A39 expression to both immunoblotting and immunohistochemical analysis of a number of glioma types and found that expression significantly varied between tumours, and all models derived from them (Fig. 1K and L and S2E).

### SLC25A39 deficiency triggers mitochondrial hyperfusion and oxidative damage

As our investigation of expression of SLC25A39 showed that it was variably expressed in GBM, and its expression was induced in hypoxia, we wanted to further investigate what downstream phenotypes this might elicit. SLC25A39 is localised to the inner mitochondrial membrane, and regulates a significant proportion of mitochondrial glutathione (mGSH) transport. We therefore hypothesised that in common with previous reports, its loss would impact on the ability of GBM cells to respond to oxidative stress. We generated multiple models of SLC25A39 deficiency (Fig. S3A) in both GBM and HAP1 cell backgrounds, and confirmed that this resulted in significant reduced mGSH (Fig. 2A).

**Figure 2:**
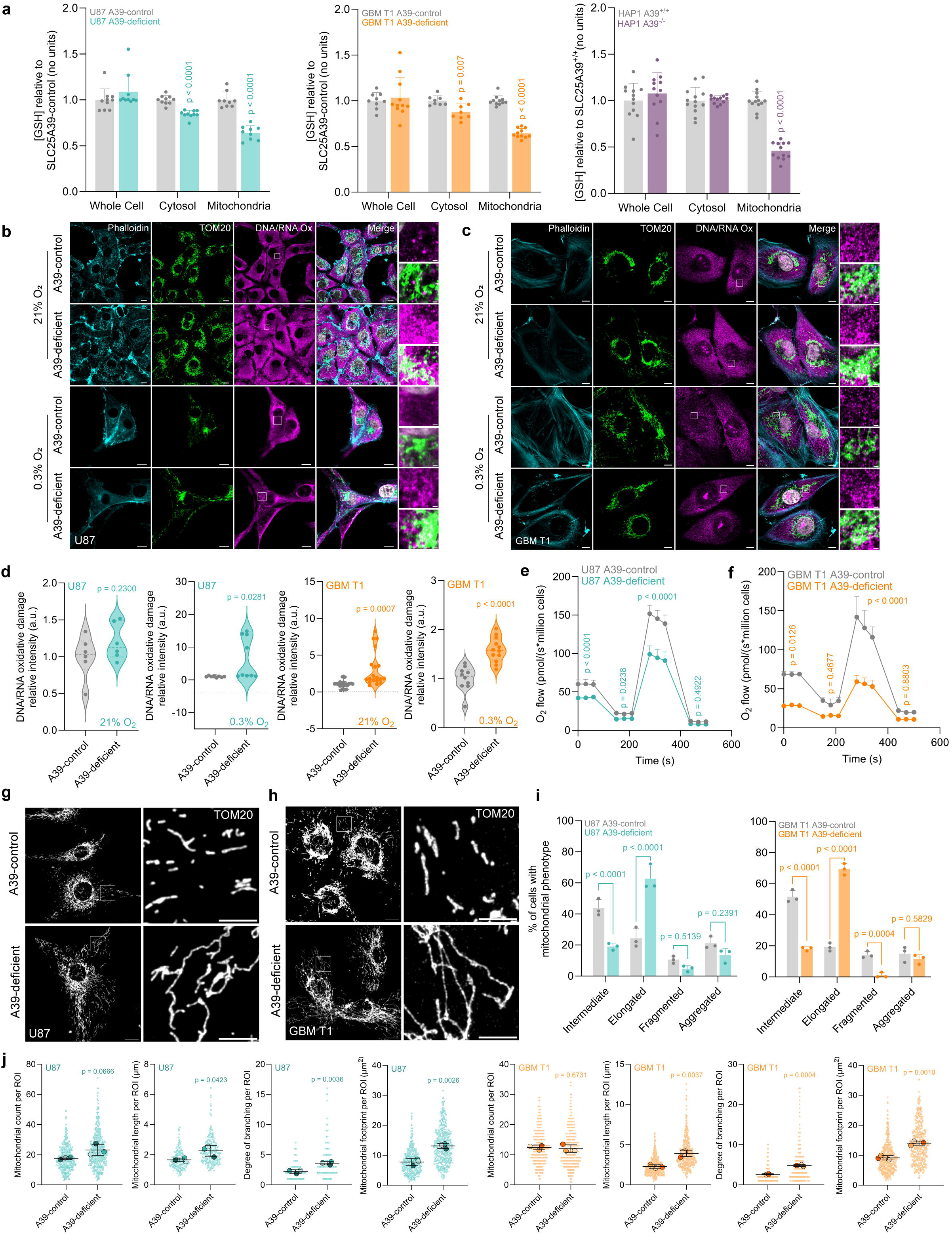
SLC25A39 depletion is associated with a hyperfused mitochondrial network and increased oxidative damage. a) Relative GSH concentration within whole cell, cytosol and mitochondria compartments in U87, GBM T1 and HAP1 SLC25A39-deficient cells relative to SLC25A39-control cells. b-c) Representative confocal images showing DNA/RNA oxidation in SLC25A39-control and -deficient U87 (b) and GBM T1 (c) cells following 24 hours incubation in 21% or 0.3% O_2_. Scale bars: cell image = 10 µm, ROI = 1 µm. d) Quantitative analysis of DNA/RNA oxidation intensity co-localised with the mitochondria (TOM20). Dashed line indicates the median; dotted lines represent upper and lower quartiles. Each data point represents an individual image. e-f) Oxygen flux plotted as a function of time. Data are presented as means; error bars, SEM. g-h) Representative confocal images of SLC25A39-control and -deficient U87 (g) and GBM T1 (h) mitochondria. Scale bars: cell image = 15 µm, ROI = 5 µm. i) Quantification of mitochondrial phenotype from (g) and (h). j) Quantification of mitochondrial parameters per ROI (225 µm^2^) from (g) and (h). Data are means; error bars, SD unless otherwise indicated; *n* = 3 independent experiments except (**a**) *n =* 2-3 independent experiments. Significance was assessed by (**a**): Mann-Whitney (two-tailed); (**d**): unpaired t-test (two-tailed); (**e-f**): one-way ANOVA with Šídák’s multiple comparisons test (using averages of 3 time points within 60 second windows; 0-60; 150-210, 280-340 and 440-500 s); (**i**): two-way ANOVA with Šídák’s multiple comparisons test; (**j**): nested t-test (two-tailed). Individual *p*-values are indicated on the respective graphs.

Using these models, we investigated oxidised DNA and RNA species as markers of downstream oxidative damage, and found that SLC25A39-deficient cells demonstrated significantly increased staining, further exacerbated in hypoxia (Fig. 2B-D and S3B and C). The increased mitochondrial-localised oxidative damage observed in SLC25A39-deficient cells raised the possibility of downstream effects on mitochondrial respiratory function. We found that loss of SLC25A39 resulted in a substantial reduction in oxygen consumption rate (OCR) indicting a clear respiratory defect (Fig. 2E and F, S3D). It has previously been shown that mitochondrial damage results in changes to the architecture of the network. We therefore performed high resolution imaging of SLC25A39 deficient cells compared to their controls, and observed a significant change in morphology, with a tubular mitochondrial structure being replaced with a hyperfused network in SLC25A39-deficient cells (Fig. 2G-J and S3E). Previous studies have demonstrated that an increased GSSG/GSH ratio —a hallmark of mitochondrial oxidative stress—strongly promotes mitochondrial fusion as a transient protective mechanism to facilitate the exchange of metabolites and to counteract localised damage^23, 24^. It is therefore likely that SLC25A39-deficient cells undergo structural remodelling as a direct response to the significant redox imbalance caused by mGSH depletion.

### SLC25A39 loss triggers significant metabolic reprogramming

To elucidate how the mitochondrial dysfunction downstream of SLC25A39 loss reshapes the wider metabolic phenotype in GBM, we performed unbiased polar metabolomics on SLC25A39-deficient U87 and patient-derived GBM T1 cells as well as the HAP1 cell model as comparator. This showed a significant metabolic divergence between control and deficient cells, characterised by significant alterations in the TCA cycle, glycolytic intermediates, and lipid precursors (Fig. 3A-C and S4A-D), all of which are consistent with being downstream of the observed mitochondrial respiratory defect. Additionally, we observed a marked decrease in the mitochondrial NAD^+^/NADH ratio in U87 and HAP1 SLC25A39-deficient cells, but a surprisingly lack of deficit in GBM T1 cell model (Fig. 3D and S4E), which overall is also consistent with the respiratory phenotype observed after loss of SLC25A39.

**Figure 3:**
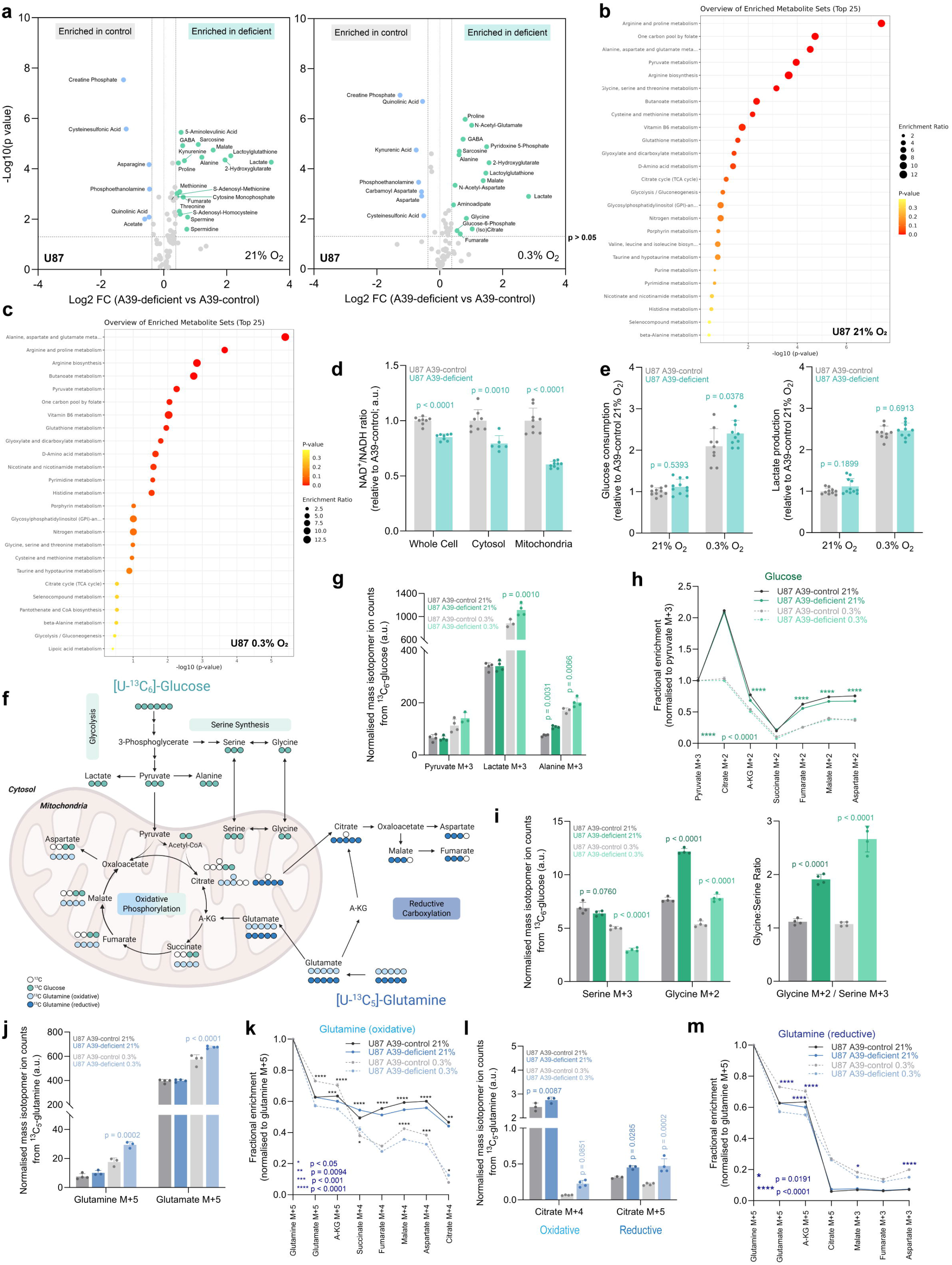
Loss of SLC25A39 alters TCA metabolite pools and shifts carbon substrate utilisation. a) Differential metabolite abundance in SLC25A39-deficient U87 cells relative to SLC25A39-control cells after 24 hours at the indicated O_2_ tensions. Significantly increased or decreased metabolites are denoted in green or blue respectively (*p* <0.05 and |log₂FC| > 1.3), non-significant in grey. b-c) Metabolite set enrichment analysis of SLC25A39-control and -deficient U87 cells under 24 hours normoxia (21% O_2_; **b**) or hypoxia (0.3% O_2_; **c**). d) NAD^+^/NADH ratio in whole cell, cytosolic and mitochondrial compartments of SLC25A39-deficient relative to -control U87 cells. e) Relative glucose consumption and lactate production after 24 hours at 21% or 0.3% O_2_. f) Schematic illustrating stable isotope labelling patterns using ^13^C_6_-Glucose (green) or ^13^C_5_-Glutamine (oxidative: light blue; reductive: dark blue). Created in BioRender (https://BioRender.com/xez62pj). g) Normalised mass isotopomer ion counts of pyruvate, lactate and alanine from 24 hours ^13^C_6_-Glucose. h) Fractional enrichments of TCA metabolites after 24 hours ^13^C_6_-Glucose (M+2 normalised to M+3 pyruvate). Data are means; error bars, SEM. i) Normalised mass isotopomer ion counts of serine and glycine and glycine:serine ratio from 24 hours ^13^C_6_-Glucose. j) Normalised mass isotopomer ion counts of glutamine and glutamate from 24 hours ^13^C_5_-Glutamine. k) Fractional enrichments of TCA metabolites after 24 hours ^13^C_5_-Glutamine (oxidative: M+5, M+4). Data are means; error bars, SEM. l) Normalised mass isotopomer ion counts of citrate from 24 hours ^13^C_5_-Glutamine (oxidative, M+4; reductive, M+5). m) Fractional enrichments of TCA metabolites after 24 hours ^13^C_5_-Glutamine (reductive: M+5, M+3). Data are means; error bars, SEM. Data are means; error bars, SD unless otherwise indicated; *n* = 4-5 independent cultures (**a-c)** or 2-3 independent experiments (**d, e**); (**g-m)** is representative from 3 independent experiments. Significance was assessed by (**a**): Student’s t-test (p <0.05, |FC| >1.3); (**d**): unpaired t-test (two-tailed); (**e**): two-way ANOVA with Šídák’s multiple comparisons test; (**g-m**) one-way ANOVA with Šídák’s multiple comparisons test (h, k, l: 21%-21%, 0.3%-0.3%). Individual *p*-values are indicated on the respective graphs.

In response to this mitochondrial defect, SLC25A39-deficient cells exhibited a compensatory shift toward aerobic glycolysis, again accentuated in hypoxic conditions, characterised by higher rates of glucose consumption and lactate secretion in the two cell models with changes in NAD^+^:NADH, but not the GBM T1 cell model (Fig. 3E and S4F). To further examine this phenotype, we traced glucose metabolism using ^13^C-[U]-glucose (Figure 3F). We noted an increase in glycolytic rate, indicated by increased glucose-derived lactate (Fig. 3G and S5A), and the expected decrease in oxidative glucose entry to the TCA cycle in hypoxia (Fig. 3H and S5B). We also observed a significant increase in the labelling of serine and glycine from glucose downstream of SLC25A39-deficiency (Fig. 3I and S5C), indicating increased serine/glycine synthesis, which may be a survival mechanism to provide the precursors for additional glutathione synthesis.

Given the surprising lack of TCA cycle defect noted from glucose labelling, we examined metabolism of the other major precursor in anabolic cells, glutamine. ^13^C-[U]-glutamine tracing showed that SLC25A39 deficiency enhanced glutamine uptake and conversion to glutamate (Fig. 3J and S5D). Examining the fractional incorporation of glutamine into downstream oxidative metabolites however suggested only a small change in contribution of glutamine to the overall pools of TCA cycle metabolites (Fig. 3K and S5E). However, the relative amount of glutamine oxidatively metabolised (e.g. citrate M+4 isotopologue) was increased in SLC25A39 cells in normoxia compared to the incorporation of glucose (Fig. 3L and S5F and G). Further differences were observed in reductive metabolism of glutamine; while relative incorporation was relative unchanged by SLC25A39 deficiency (Fig. 3M and S5H), change in the pool size meant that increased glutamine reduction to citrate was observed (Fig. 3L and S5G). Overall, while glucose metabolism in SLC25A39 deficient mitochondria was unchanged, glutamine catabolism was increased in both normoxia and hypoxia.

Our data showed that SLC25A39-deficient cells have striking levels of oxidative stress and hyperfused mitochondria. Previous studies have suggested that oxidative stress drives changes in lipid metabolism, some of which can be induced by changes in NADPH availability, which is required for both the antioxidant response and lipid biosynthesis^25, 26^. The NADP^+^:NADPH ratio was therefore examined, and surprisingly no consistent alterations observed between the cell models (Fig. S6A). However, profiling of the fatty acids in both GBM models showed a consistent increase in the fatty acid desaturation index after loss of SLC25A39, with both MUFAs and PUFAs total abundance or class significantly increased even though total fatty acid abundance remained unchanged (Fig. 4A and S6B-D). Given the propensity of unsaturated fatty acids for peroxidation by ROS, we assessed this by quantifying malondialdehyde (MDA), and noted a significant increase (Fig. 4B), consistent with increased lipid oxidation. Under these conditions, unsaturated fatty acids are often sequestered into lipid droplets as a protective mechanism. BODIPY staining of the cell models showed that loss of SLC25A39 increased lipid droplet size or number, or both (Fig. 4C-E).

**Figure 4:**
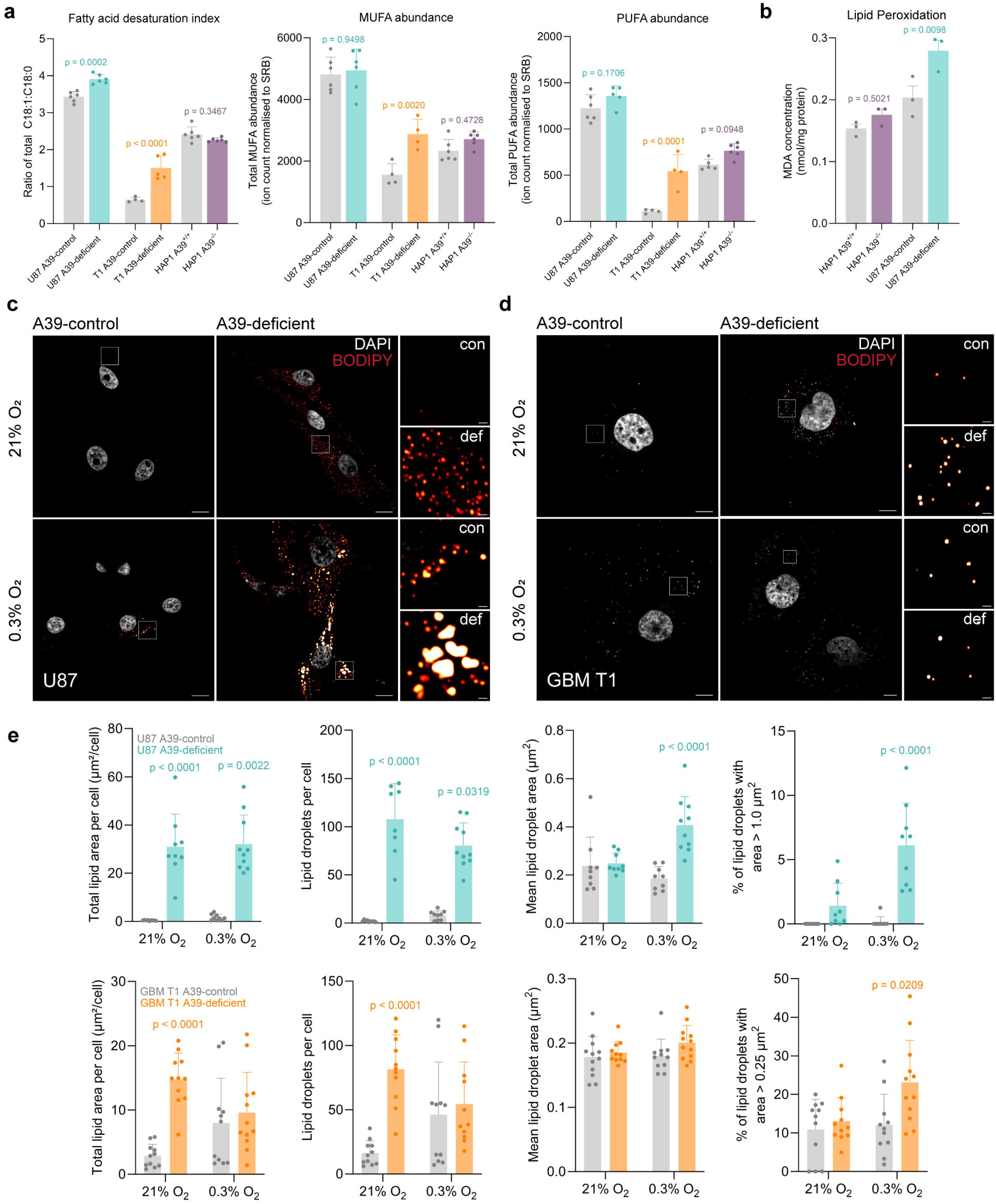
SLC25A39 loss increases lipid droplet burden and oxidative lipid damage. a) Fatty acid desaturation index (calculated as the ratio of total monounsaturated C18:1 species to total saturated C18:0 species) and total monounsaturated (MUFA) and polyunsaturated (PUFA) fatty acid abundance in U87 and GBM T1 SLC25A39-control and-deficient and HAP1 SLC25A39^+/+^ and SLC25A39^-/-^ cells. b) Cellular lipid peroxidation assessed by malondialdehyde (MDA) concentration in HAP1 SLC25A39^+/+^ and SLC25A39^-/-^ and U87 SLC25A39-control and -deficient cells. c-d) Representative confocal images of neutral lipid droplets stained with BODIPY 493/503 in U87 (**c**) and GBM T1 (**d**) SLC25A39-control (con) and -deficient (def) cells after 24 hours incubation at indicated oxygen tension. Nuclei represented in grey, lipid droplets using Sun LUT. Cell image scale bar = 10 µm, ROI = 1 µm. e) Quantitative analysis of lipid droplet characteristics in (**c**) and (**d**). Data are means; error bars, SD; *n* = 3-6 independent cultures (**a**) or 3 independent experiments (**b-e**). Significance was assessed by (**a**, **b**): one-way ANOVA with Šídák’s multiple comparisons test; (**e**): Kruskal-Wallis test with Dunn’s multiple comparisons test. Individual *p*-values are indicated on the respective graphs.

### SLC25A39 deficiency blunts GBM cell growth

The mitochondrial and metabolic defects observed following SLC25A39 loss led us to evaluate the functional impact on GBM cell growth. In standard 2D culture, SLC25A39 deficiency in both U87 and the treatment-resistant GBM T1 line resulted in a significant reduction in cell proliferation rates under both normoxic and hypoxic conditions (Fig. 5A, B and S7A). To better simulate the complex physiological architecture and metabolic gradients of a tumour mass, we utilised 3D spheroid models. Consistent with the 2D findings, U87 and GBM T1 SLC25A39-deficient cells formed spheroids that grew significantly slower compared to WT controls (Fig. 5C and D, S7B-D).

**Figure 5:**
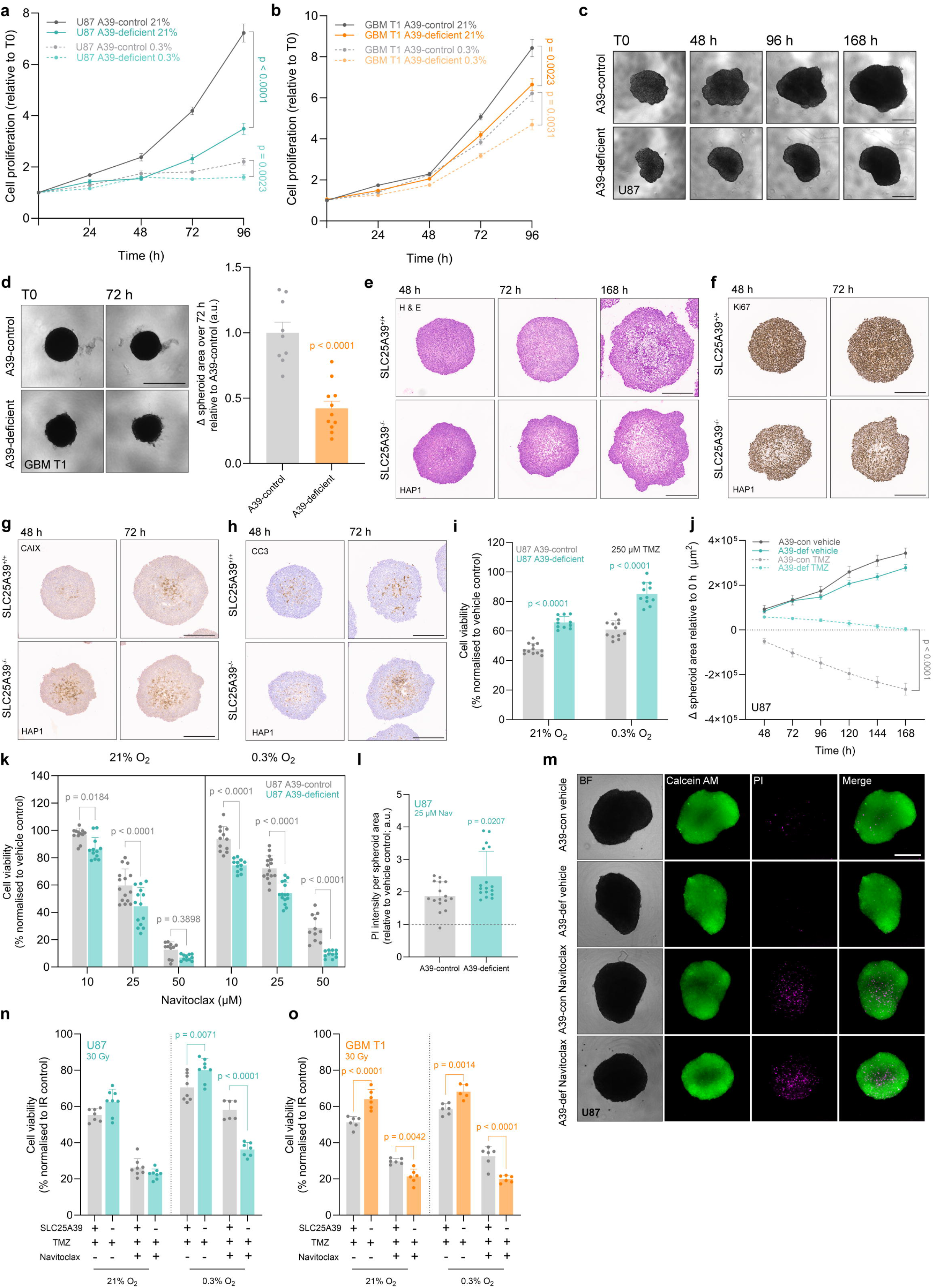
SLC25A39 depletion impairs GBM cell growth and modulates sensitivity to therapeutic stress. a-b) Proliferation of SLC25A39-control and -deficient U87 (**a**) and GBM T1 (**b**) cells at 21% or 0.3% O_2_, relative to time zero (T0). Error bars ± SEM. c-d) Representative brightfield images of SLC25A39-control and -deficient U87 (**c**) and GBM T1 (**d**). Scale bar = 500 µm. Quantification in (**d**) shows change in spheroid area (µm^2^) over 72 hours of SLC25A39-deficient GBM T1 cells relative to T0 and SLC25A39-control. e-h) Immunohistochemistry of HAP1 SLC25A39^+/+^ and SLC25A39^-/-^ spheroids at the indicated time points. Cell distribution and spheroid integrity is shown using H & E (**e**); proliferative capacity (Ki67; **f**); hypoxia (CAIX; **g**); apoptosis (cleaved caspase-3; **h**). Scale bar = 300 µm. i) Viability of SLC25A39-control and -deficient U87 cells after 250 µM temozolomide (TMZ) for 72 hours, relative to vehicle at the indicated O_2_ tensions. j) Change in spheroid area (µm^2^) of vehicle-or 500 µM TMZ-treated SLC25A39-control and -deficient U87 spheroids relative to the start area (T0) for each spheroid. Error bars ± SEM. k) Viability of SLC25A39-control and -deficient U87 cells treated with the indicated navitoclax (Nav) concentrations for 24 hours. l) Propidium iodide (PI) staining intensity of vehicle-or 25 µM navitoclax-treated SLC25A39-control and -deficient U87 spheroids, normalised to spheroid area and relative to vehicle. m) Representative images showing viable (calcein AM; green) and dying/dead (PI; magenta) U87 spheroids after 25 µM navitoclax for 96 hours. Scale bar = 500 µm. n-o) Viability of U87 (**n**) and GBM T1 (**o**) cells 48 hours after 30 Gy irradiation, treated with 250 µM (U87) or 500 µM (GBM T1) TMZ alone, or combined with 10 µM navitoclax, relative to irradiation only at the indicated O_2_ tensions. Data are means; error bars, SD unless indicated. *n* = 3-4 independent experiments, except **e-h** which are representative images from 96 spheroids per condition. Significance was assessed by (**a, b, d**): unpaired t-test (two-tailed); (**i-k**): two-way ANOVA with Tukey’s multiple comparison test; (**l**): Mann-Whitney (two-tailed); (**n, o**): one-way ANOVA with Šídák’s multiple comparisons test. *p*-values are indicated on the respective graphs.

To provide insights into the basis of the growth defect, we performed immunohistochemical staining of sections of the 3D structure, noting both a reduction in sphericity and decreased proliferation (Ki67+ cells; Fig. 5E and F). Interestingly, we also noted changes in spheroid hypoxic response (carbonic anhydrase IX, CAIX; Fig. 5G) and apoptotic cell death (cleaved caspase-3; Fig. 5H). Given that SLC25A39 impairs oxygen consumption (Fig. 2E and F, S3D), but increases oxidative damage (Fig. 2B and C, S3B), it is likely that HIF-1 stabilisation leading to CAIX upregulation is through the well-described ROS-mediated route^27, 28^. This suggests that ROS generation downstream of SLC25A39 deficiency impairs proliferation and leads to increased cell death.

### SLC25A39 expression alters therapeutic response and provides a targetable vulnerability in TMZ-resistant GBM

As expression of SLC25A39 varies between and within GBM tumours, this could represent a therapeutic opportunity for these incurable cancers. We first examined whether loss of SLC25A39 altered the response to standard of care chemotherapeutic agent, TMZ. Interestingly, SLC25A39-deficiency increased resistance to TMZ across all cell models (Fig. 5I, J and S7E-J), suggesting that tumours with low SLC25A39 expression may be more resistant to this therapy. To identify other therapeutic options that instead synergise with loss of SLC25A39, we turned to those that directly target mitochondria. We first examined navitoclax (ABT-263), a potent BH3 mimetic that inhibits Bcl-2/Bcl-xL, thereby inducing apoptosis. Strikingly, SLC25A39-deficient cells exhibited significantly increased sensitivity to navitoclax, particularly under hypoxic conditions (Fig. 5K and S7K and L). Consistent with this, SLC25A39-deficient 3D spheroid models also show increased sensitivity to cell death (Fig. 5L and M, S8A-E). This sensitisation was also observed with a similar BH3 mimetic – ABT-737 – which led to significant cell death and loss of spheroid integrity (Fig. S8F-H).

Finally, we explored whether combining navitoclax alongside TMZ and ionising radiation (IR) – the other treatment modality used as standard of care for GBM – could alleviate the resistance to TMZ observed downstream of SLC25A39 deficiency. We observed that navitoclax significantly increased cell killing in combination with IR and TMZ (Fig. 5N and O), suggesting that tumours with significant intratumoural heterogeneity in SLC25A39 expression may benefit from the addition of this agent.

## Discussion

Since its identification in 2021, SLC25A39 has been shown to be the mitochondrial glutathione transporter whose expression is regulated by cell stress and its environment, and is therefore critical for maintaining mitochondrial health. Our data demonstrate that SLC25A39 is a HIF-target gene whose expression is induced by hypoxic conditions that characterise GBM. Indeed, we also show that SLC25A39 expression varies within and between GBM tumours, and while mRNA correlates with tumoural hypoxia, the complexity of the regulation of protein stability by multiple metabolic stimuli show that assessment of protein expression is critical to understand the implications of differential expression on tumour biology.

Our data suggest that in vitro, loss of SLC25A39 led to downstream effects on both mitochondrial phenotype and function, including the evolution of a hyperfused mitochondrial network. This is consistent with previous work showing that decreased mitochondrial GSH:GSSG ratio alters oxidises key residues in mitofusin, leading to its oligomerisation and mitochondrial fusion. This may therefore be a protective response to avoid local depletion of GSH, but in a situation where glutathione transport into the mitochondrion itself is deficient, it likely maladaptive.

We documented significant downstream metabolic reprogramming after loss of SLC25A39. While some cell models showed a more significant variation in aspects of redox control and glycolysis, the data across multiple pathways suggest that all models reflected different metabolic responses deployed to maintain biosynthetic and redox capacity downstream of mitochondrial dysfunction. Interestingly, one consistent response was towards serine/glycine synthesis, which might be expected to contribute to de novo nucleotide biosynthesis and glutathione biosynthesis.

GBM are highly therapy resistant cancers. We therefore examined whether alterations in mitochondrial glutathione and the oxidative stress arising from it might alter response to conventional therapy. Interestingly, we showed that SLC25A39 deficiency leads to increased resistance to TMZ – an alkylating agent still used as first line therapy in patients. It is possible that the metabolic changes we observed could underpin this, as enhanced nucleotide synthesis could permit a more efficient DNA damage repair mechanism. However, loss of SLC25A39 sensitised GBM cells to agents targeting mitochondrial apoptosis pathways, such as navitoclax. This evidence is consistent with other studies suggesting that GBM stem cells are dependent on Bcl-xL and MCL1 for survival^29, 30^, and our data suggest that loss of SLC25A39 primes this further.

Our findings therefore position SLC25A39 as a GBM-relevant determinant of mitochondrial redox homeostasis and therapy responsiveness. We showed that SLC25A39 expression varies both within and between GBM tumours, with some samples showing no detectable expression. This is consistent with emerging evidence that SLC25A39 status may differ across cancer types, with coinciding diagnostic and prognostic value^31^. Recently, Yeh et al.,^32^ showed that SLC25A39-dependent mitochondrial glutathione import supports metastatic colonisation in breast cancer via ATF4 signalling, however loss of SLC25A39 does not fully abrogate metastasis, implying existence of compensatory mechanisms. We similarly observed no detectable SLC25A39 expression in a small number of recurrent GBM tissue samples raising the possibility that SLC25A39 may dispensable or selected against during disease progression in some tumours. If and/or when SLC25A39 expression changes during GBM evolution and how that relates to environmental triggers and treatment response, therefore warrants further investigation in larger longitudinal cohorts.

From a therapeutic perspective, our data suggest that SLC25A39 could serve as a stratification marker for treatment response in GBM. SLC25A39-deficient cells showed in vitro, increased resistance to TMZ but a greater dependence on mitochondrial redox buffering — a vulnerability we propose may be exploitable through combination of therapeutic agents that target mitochondrial stress with current GBM standard of care to target the most therapy resistant cells in these tumours and thereby potentially increase patient survival.

## Acknowledgements

We would like to thank and acknowledge Mr. Robert Spier whose generous donation supported I.R. The authors would also like to acknowledge Jamie R.M. Webster for training and technical assistance, the Light Microscopy Facility at the MRC MBU in Cambridge, Genomics Birmingham for access to the irradiator platform, the University of Birmingham Imaging Facility for access to microscopy platforms, and also thank the Metabolic Tracer Analysis Core at the University of Birmingham for sample analysis and technical assistance. We would also like to thank the patients who generously provided specimens used in this research.

## Author Contributions

D.A.T and K.L.E conceptualised the study. Investigation, validation and methodology was carried out by K.L.E., Y.L, Y.F, I.R., S.R.C., B.M., H.P-C., C.N. and J.R. Formal analysis was performed by K.L.E, Y.L, Y.F, S.R.C, B.M, J.R and J.P, while K.L.E, Y.L, S.R.C, C.N, V.W, I.R, B.M, J.R and Y.F undertook data curation. K.L.E, D.A.T, J.P and W.H were responsible for funding acquisition and project administration. Resources were made available by V.W., J.P., W.H. and D.A.T. Writing was undertaken by K.L.E, Y.L. and D.A.T, while reviewing and editing was performed by all authors.

## Methods

### Cell culture and model generation

Human GBM lines were obtained from the American Type Culture Collection (ATCC; T98G, CRL-1690) and CLS Cell Lines Services GmbH (Germany; A172 #300108, U343 #300365, U87 #300367). HAP1 SLC25A39^-/-^ (knockout, #HZGHC002113c010) cells were obtained along with the accompanying HAP1 parental line (SLC25A39^+/+^ wildtype, #C631) from Horizon Discovery (UK).

GBM T1 cells (Pt 1) are a patient cell line derived from a therapy-resistant section of GBM tissue, as described in Eales *et al*^33^. Additional GBM patient lines (Pt 2 and Pt 3) were generated as outlined in Eales *et al*^33^, as part of the Brain Surgical Tissue for Advanced Tumour Models (BRAINSTAT; REC Reference No. 20/EM/0211) from consented patients undergoing their first maximal resection at the Queen Elizabeth Hospital Birmingham, UK. All patients provided full, informed, written consent in accordance the Declaration of Helsinki and procedures were conducted in accordance with good clinical practice (GCP) and ethical standards outlined in The Human Tissue Act 2004.

T98G, A172, U343, U87, GBM T1 and patient-derived cells were cultured in antibiotic-free Dulbecco’s Modified Eagle’s Medium (DMEM) high glucose (Gibco, 41965039) and HAP1 cells in antibiotic-free Iscove’s Modified Dulbecco’s Medium (IMDM; Gibco, 12440053 and 21980-032). All cells were cultured with addition of 10% foetal bovine serum (FBS; Sigma-Aldrich) and maintained under standard culture conditions (humidified incubator 37°C, 5% CO_2_). For metabolic experiments, dialysed FBS was supplemented instead, generated in house using 3.5kDa MWCO Slide-A-Lyzer™ Dialysis Flasks (Thermo, 13494199). For hypoxia studies, cells were plated under standard conditions and left to adhere, before being transferred to a Whitley H35 Hypoxystation (Don Whitley Scientific, UK) set to 0.3% O_2_ (37 °C, 5% CO_2_) for the indicated experimental time. All cell lines were subject to routine mycoplasma testing (Geneflow, K1-0210), with no positive results returned. Cell lines were successfully authenticated using STR profiling services provided by NorthGene™ (Biofortuna Limited, UK).

#### SLC25A39 knockdown generation

Stable SLC25A39 control (A39-control) or knockdown (A39-deficient) U87 and GBM T1 cell models were generated using the lentiCRISPR v2 CRISPR-Cas9 system. Three candidate SLC25A39-targeting single-guide RNAs (sgRNAs) were constructed with the following 20-nucleotide spacer sequences: sgHsA39-1, 5′-CTAGGTCTTACTGAAGAGAG-3′; sgHsA39-2, 5′-GGACACCATTGCAATACAGG-3′; and sgHsA39-3, 5′-TGACCTCTACGCACCCATGG-3′. The non-targeting control sgRNA (sgNT) contained the spacer 5′-AAATGTGAGATCAGAGTAAT-3′. Guide oligonucleotides were synthesised by Sigma-Aldrich/Merck Life Science UK as 200-nmol desalted DNA oligonucleotides. Complementary oligonucleotides were annealed and cloned into BsmBI-digested lentiCRISPR v2 (a gift from Feng Zhang; Addgene plasmid #52961; RRID:Addgene_52961)^34^. Correct plasmid construction was confirmed by commercial whole-plasmid sequencing (Plasmidsaurus).

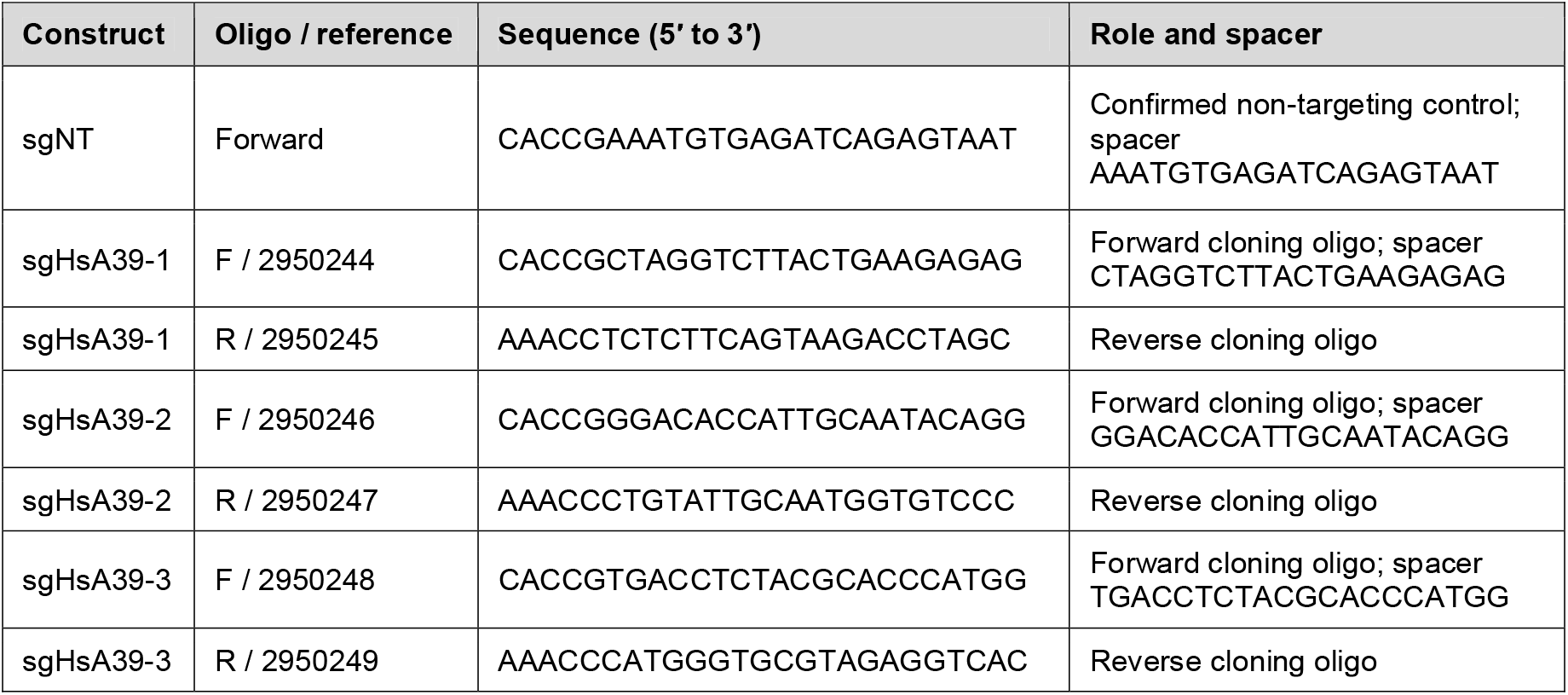

Lentiviral particles were produced in HEK293T cells by co-transfecting each lentiCRISPR v2 transfer construct with the second-generation packaging plasmid psPAX2 and the VSV-G envelope plasmid pMD2.G (gifts from Didier Trono; Addgene plasmid # 12260; RRID:Addgene_12260 and #12259; RRID:Addgene_12259 respectively)). For each well of a 6-well plate, 1 µg transfer plasmid was combined with 1 µg of the packaging/envelope-plasmid mixture, in which psPAX2 and pMD2.G were present at a 3:1 mass ratio (2 µg total DNA per well). Plasmid DNA was transfected using 4 µL DharmaFECT kb DNA Transfection Reagent (Horizon Discovery). Viral supernatants were collected 48 h after transfection and passed through a 0.45-µm filter. For transduction, filtered viral supernatant was mixed 1:1 (v/v) with complete growth medium and applied to U87 or GBM T1 cells in the presence of 10 µg/mL polybrene. Transduced cells were selected with 1 µg/mL puromycin and were maintained as pooled populations rather than single-cell-derived clones. SLC25A39 depletion was confirmed by immunoblotting before experimental use and guide Sg-2 was selected for experiments.

### siRNA transfection

For transient knockdown of HIF-1α and -2α expression, cells were transfected according to manufacturer’s instructions using either 25 nM ON-TARGETplus non-targeting control pool siRNA (siNT; D-001810-10) or ON-TARGETplus Human *HIF1A* (SMARTPool L-004018-00) or ON-TARGETplus Human *EPAS1* (HIF-2α; SMARTPool L-004814-00) with DharmaFECT 1 transfection reagent (T-2001), all from Horizon Discovery. Cells were cultured under standard conditions for 48 h to ensure sufficient silencing before being transferred to 0.3% O_2_ for the indicated time.

### Western blotting

Cells were washed with ice-cold 1x PBS and protein lysates were lysed using 1 x Laemmli buffer (Sigma-Aldrich, S3401-1VL). Samples were heated to 95 °C for 10 min followed by loading of equal amounts of sample onto 10% Tris-Glycine polyacrylamide gels and resolved by sodium dodecyl sulphate polyacrylamide gel electrophoresis (SDS-PAGE) using the mini-PROTEAN Tetra system (Bio-Rad). Proteins were then transferred to nitrocellulose membranes (0.2 µm pore; IB23001, Invitrogen) using the iBlot™ gel transfer system (Invitrogen). Depending on antibody specifications, membranes were either blocked in PBS-0.1% Tween (PBS-T) containing 5% bovine serum albumin (BSA; Sigma-Aldrich A7906) or 5% non-fat dried milk powder (Marvel) for 1 h at room temperature (RT). After blocking, membranes were cut and incubated with primary antibodies overnight at 4 °C. The next morning, membranes were washed with PBS-T and incubated with the appropriate horseradish-peroxidase-conjugated secondary antibodies (Cell Signaling, mouse #7076 or rabbit #7074) for 1 h at RT. Membranes were then washed and visualised using Clarity™ Western ECL Substrate (Bio-Rad, #1705060) and imaged using the Bio-Rad ChemiDoc XRS+ imaging system and Image Lab 6.0.1 software. Primary antibodies used were as follows: anti-β-tubulin (1:2000 in 5% BSA; Cell Signaling, #2128), anti-HIF-1α (1:500, 1% milk; BD Transduction, 610959), anti-HIF-2α (1:500 in 1% milk; Novus Biologicals, #NB100-122), anti-SLC25A39 (1:1000, 5% BSA; Proteintech, 14963-1-AP) and anti-TOM20 (1:2000, 5% milk; Abcam, ab78547).

### Determining SLC25A39 expression in patient tissue slices

#### Clinical Sample collection

Formalin-fixed, paraffin-embedded (FFPE) surgical specimens were collected from 23 brain tumour patients (20 glioblastomas, 2 low grade gliomas and 1 astrocytoma) treated at the Department of Neurosurgery, Huashan Hospital, Fudan University, Shanghai. Ethical approval for this study was granted by the Institutional Review Board of Huashan Hospital (Approval No. KY2022-496), with signed informed consent secured from all patients. Histopathological diagnoses were independently verified by two pathologists.

#### Immunohistochemistry (IHC)

IHC was conducted according to the manufacturer’s protocol. Briefly, sectioned FFPE tissues underwent deparaffinisation, heat-induced epitope retrieval, and serum blocking before being probed with a rabbit anti-SLC25A39 monoclonal primary antibody (1:100 dilution; Cell Signaling Technology, Cat #33871T). Signal amplification was performed using horseradish peroxidase (HRP)-linked secondary antibodies (Vector Laboratories, Cat #MP-7500), followed by chromogenic visualisation with 3,3′-diaminobenzidine (DAB) substrate (Servicebio, Cat #C0105S). Slides were counterstained with haematoxylin and digitised using an Olympus VS200 slide scanner. Staining intensity was classified as negative if no specific cytoplasmic expression was detected; weak positive for faint or partial granular staining in the cytoplasm; and medium-strong positive when tumour cells displayed distinct, prominent, and intense dark staining.

#### Multiplex immunofluorescence

Staining was performed using a TSA-based multiplex immunofluorescence assay kit (Ruchuang, Cat#RC0086Plus-34 R) according to the manufacturer’s instructions. Briefly, FFPE tissue sections were deparaffinised, rehydrated, subjected to heat-induced antigen retrieval, and blocked before sequential rounds of immunostaining. In each staining cycle, sections were incubated with the corresponding primary antibody, followed by an HRP-conjugated secondary antibody (Vector Laboratories, Cat#MP-7500) and tyramide signal amplification using the appropriate Opal fluorophore working solution for 10 min. After each round of TSA labelling, the antibody complexes were removed by heat-mediated stripping before initiation of the subsequent staining cycle. Primary antibodies included anti-HIF-1α (1:200; Proteintech, Cat#66730-1-Ig) and anti-SLC25A39 (1:200; Cell Signaling Technology, Cat#33871T). After completion of all staining cycles, nuclei were counterstained with DAPI. Whole-slide fluorescence images were acquired using an Olympus VS200 slide scanner. *Quantification*: From 3 patients, 4 spatially independent HIF-1α-positive regions and 4 spatially independent HIF-1α-negative regions were selected based on the presence or absence of detectable HIF-1α signal. Images used for comparison were acquired under comparable imaging conditions. Fluorescence intensity was quantified in Fiji/ImageJ.

### Bioinformatical analysis

#### TCGA/GDC and GLASS cohorts and transcriptional subtype classification

STAR-derived gene-expression data, somatic mutation profiles, and clinical annotations for TCGA-GBM were obtained from the Genomic Data Commons and cBioPortal. Cases carrying IDH mutations were excluded to establish an operationally defined IDH-wild-type cohort. Gene-expression and clinical data from the Glioma Longitudinal Analysis Consortium (GLASS) cohort were additionally obtained from cBioPortal. Proneural, Classical, and Mesenchymal subtypes were assigned using the ssgsea.GBM.classification Rpackage (https://github.com/zhaoliang0302/ssgsea.GBM.classification). Each sample was assigned to the subtype with the smallest permutation-derived *p* value.

#### Public single-cell RNA-sequencing data and hypoxia classification

Single-cell RNA-sequencing data were obtained from the GBmap glioblastoma dataset (<u>reference atla2</u>). The atlas-provided raw RNA counts, log-normalised expression values, UMAP coordinates, and cell-type annotations were used. Cellular hypoxia status was assigned using the cellular hypoxia predicting framework (CHPF) 3, applied separately to each study (https://github.com/yihan1221/CHPF).

#### Bulk RNA expression, survival, and enrichment analyses

SLC25A39 expression was compared across transcriptional subtypes and histological grades using two-sided Wilcoxon rank-sum tests. Overall survival (OS) and progression-free interval (PFI) were analysed after excluding cases with missing time or event data. Kaplan-Meier curves were compared using two-sided log-rank tests. For genome-wide comparisons, samples were divided at the cohort median of SLC25A39 expression. Genes were tested using two-sided Wilcoxon rank-sum tests with Benjamini-Hochberg correction. Pathway activities were derived from the Molecular Signatures Database (MSigDB) and quantified using GSVA with a Gaussian kernel. Spearman correlations between SLC25A39 expression and pathway scores were calculated within each transcriptional subtype. Correlations with individual pathway genes were adjusted using the Benjamini-Hochberg procedure.

### Immunocytochemistry and Confocal imaging

#### DNA/RNA Oxidation

Cells were seeded onto glass bottom confocal dishes (VWR, 734-2906), pre-coated with poly-L-ornithine (Sigma P3655) and fibronectin (Corning, 354008) and left to adhere overnight. The following day dishes were either left under standard conditions or transferred to 0.3% O_2_. After 24 h, cells were fixed using 10% neutral buffered formalin (Sigma-Aldrich, HT501128) for 15 min at RT, followed by quenching with 0.1 M glycine (Sigma-Aldrich) for 5 min at RT. Cells were washed with 1x PBS and then permeabilised with 0.1% Triton-X-100 Sigma-Aldrich, T8787) for 8 min, followed by a further 3 PBS washes. Cells were then blocked using 5% BSA in PBS for 30 min before being incubated with primary antibodies in 5% BSA in PBS overnight at 4 °C. The primary antibodies used were as follows: Anti-DNA/RNA Damage antibody (1:200; Abcam, ab62623), CoraLite 594-Phalloidin conjugated (1:250; Proteintech, PF00003) and TOM20 (1:500; ab78547). The following day, cells were washed 3 times with 1x PBS, before being incubated with Goat anti-Mouse IgG (H+L) Cross-Adsorbed Secondary Antibody, Alexa Fluor™ 488 (1:500; Invitrogen, A-11001) and Goat anti-Rabbit IgG (H+L) Cross-Adsorbed Secondary Antibody, Alexa Fluor™ 647 (1:500; Invitrogen, A-21244) in 5% BSA in PBS for 1 h at RT. Cells were then washed with 1 x PBS and incubated with NucBlue™ Fixed Cell ReadyProbes™ Reagent (DAPI) (Invitrogen, R37606) for 10 min at RT. Cells were then washed for a final 3 times with PBS and mounted to slides using Mowiol® 4-88 mounting medium (Sigma-Aldrich, 81381).

Cells were then imaged a ZEISS LSM 880 Axio Observer confocal microscope with Airyscan (Carl Zeiss Microscopy, Germany) equipped with an Alpha Plan-Apochromat 100x/1.46 Oil DIC M27 Elyra objective. Airyscan imaging was performed using a 1 Airy Unit pinhole, with Airy reconstruction using the ZEN 2012 SP1 software (Carl Zeiss Microscopy, Germany; black edition, version 8.1).

DNA/RNA oxidation was quantified directly from raw Zeiss ZEN CZI files using a custom Python workflow (mitoverse; https://github.com/sstoner/mitoverse/tree/master). The mitochondrial-marker channel (TOM20; channel index 0) and the DNA/RNA damage fluorescence channel (target channel index 2) were extracted as two-dimensional images. For CZI files containing additional non-spatial dimensions, the workflow selects the first plane (index 0) and does not calculate a z-projection. The TOM20 channel was segmented by Otsu thresholding to generate a binary mitochondrial mask. Small holes within the mask were filled using an area threshold of 64 pixels, followed by binary opening with a disk-shaped structuring element of 3-pixel radius to reduce isolated noise. The resulting mitochondrial mask was applied to the corresponding DNA/RNA damage channel. The primary plotted metric was the arithmetic mean of the DNA/RNA damage pixel intensities within the mitochondrial mask (Average_Intensity_in_Mitochondria) for each image; total DNA/RNA damage intensity, mitochondrial-mask pixel count, mask coverage and mean TOM20 intensity were exported as auxiliary quality-control outputs. Each data point represents one image. Because the repository does not perform relative-intensity normalisation, exported mean intensities were normalised during graph preparation. Normalisation was performed separately within each independent experiment by dividing each value by the mean value of the corresponding same-experiment scramble-control (scr) or wild-type (WT) group, such that the matched control mean was 1.

#### Mitochondrial Morphology

Cells were seeded onto poly-L-lysine (Sigma-Aldrich, P4707) coated coverslips and left overnight. The following day, cells were fixed with 5% paraformaldehyde (PFA) in PBS at 37 °C for 15 min, after which cells were quenched with 50 mM ammonium chloride in PBS. Coverslips were washed three times with PBS then permeabilised in 0.1% Triton-X-100 (Sigma-Aldrich, T8787) for 8 min, followed by a further 3 PBS washes. Cells were then blocked in 10% FBS in PBS for 30 min at RT, followed by incubation with anti-TOM20 (1:1000; Proteintech, 11802-1-AP) in 5% FBS in PBS for 2 h at RT. Cells were then washed 3 times with 5% FBS in PBS and incubated with Goat anti-Rabbit IgG (H+L) Highly Cross-Adsorbed Secondary Antibody, Alexa Fluor™ 488 (1:1000; Invitrogen, A-11034). Following incubation, cells were washed 3 times in PBS and once with distilled water, before mounting onto slides using ProLong™ Diamond Antifade Mountant (Invitrogen, P36961).

Mitochondrial images were acquired on a Nikon Eclipse TiE inverted microscope equipped with a 4-channel integrated laser engine (ILE-400; Andor) and an Andor Dragonfly 500 spinning-disk confocal system, using a 100x oil-immersion objective lens (NA1.4). Z-stack images (7 slices, 0.2 µm each) were captured with a Zyla 4.2 PLUS sCMOS camera using Fusion software (Andor). Imaging parameters (including exposure time and laser intensity) were kept consistent across all images within the same experiment.

Images were first converted to TIFF format and compiled by ‘max projection’. Quantitative mitochondrial morphological analysis was performed by first randomly selecting 225 µm^2^ regions of interest (ROIs) located at the cell periphery using FIJI software^35^ (version ImageJ 1.54p) These ROIs were then analysed using the FIJI macro MitoMAPR^36^, which quantified the number, length, degree of branching (number of junction points/number of mitochondrial network in a ROI) and mitochondrial footprint (sum of the area covered by GFP signal from each mitochondrial particle in a ROI). Representative images shown have been ‘smoothed’ once using FIJI software. A minimum of 20 images were analysed from 3 independent experiments for each cell line. For mitochondrial phenotype analysis, mitochondria were classified as ‘intermediate’ (tubular and separated), ‘elongated’ (long and connected) ‘fragmented’ (short and spherical) and ‘aggregated’ (condensed to perinuclear area)^37, 38^. A minimum of 30 cells per condition from 3 independent experiments for each cell line were analysed.

#### Lipid droplet staining

Following 24 h incubation in hypoxia, cells were fixed using 10% neutral buffered formalin for 15 min at RT. Cells were then washed using 1 x PBS and permeabilised with 0.1% Triton-X-100 for 8 min at RT. Cells were washed again and incubated with 2 µM BODIPY 493/503 (Cambridge Bioscience, T36957) solution in PBS for 30 min at 37 °C. Cells were then washed twice with PBS and incubated with NucBlue™ Fixed Cell ReadyProbes™ Reagent (DAPI) (Invitrogen, R37606) for 10 min at RT. Cells were washed for a final 3 times with 1x PBS and mounted to slides using Mowiol® 4-88 mounting medium (Sigma-Aldrich, 81381). Images were acquired as outlined in the DNA/RNA oxidation section.

Lipid droplet analysis was performed using FIJI software (ImageJ). Raw BODIPY fluorescence images were processed identically across all experimental conditions. Background fluorescence was subtracted (rolling ball radius = 20 pixels), and images were thresholded using the Otsu method. Binary masks were generated and adjacent lipid droplets were separated using Watershed segmentation. Lipid droplets were identified using Analyze Particles, with a size threshold of 0.1-100 µm^2^. Total lipid droplet count, droplet area and total droplet area were quantified for each image. Nuclei were manually counted from the corresponding DAPI images and used to normalise droplet number and total lipid droplet area on a per-cell basis. Distribution analysis was performed by binning individual droplet areas and calculating the percentage of droplets within each size category for each image. A minimum of 9 images from 3 independent experiments were analysed for each cell line.

### High Resolution Respirometry

Oxygen consumption by intact cells was measured using an O2k high-resolution respirometer (Oroboros Instruments, Innsbruck, Austria) at 37 °C. Before cell loading, the instrument chambers were equilibrated for at least 2 h with DMEM + 10% FBS, and a cell-free medium blank was recorded. ROUTINE respiration was recorded after the oxygen-flux signal had stabilised. Oligomycin was then added to a final concentration of 1 µM to determine oligomycin-inhibited LEAK respiration. FCCP was subsequently added to a final chamber concentration of 2 µM to determine maximal uncoupled electron-transfer-system capacity. Rotenone and antimycin A were then added together to final concentrations of 1 µM each to inhibit complexes I and III, respectively.

Oxygen flux was normalized to 10 cells and plotted as a function of time. Each plotted time point represents the mean oxygen-flux value over an approximately 30-60s interval. Oxygen-flux values were corrected only by subtracting the cell-free medium blank recorded before cell loading; residual oxygen consumption measured after rotenone plus antimycin A was not subtracted.

### Untargeted metabolomics by triple quadrupole LC-MS (LC-QqQ)

*Extraction*: Media was removed from cells after the indicated incubation times and cells were washed once with 0.9% (w/v) NaCl in LC-MS-grade water (Biosolve, BIO-23214102-1). The plate was placed on a metal plate over dry ice and 250 µL of ice-cold extraction mix (80% LC-MS-grade methanol; Biosolve, BIO-13687802; 20% LC-MS-grade water containing the internal standard 0.25 µg/mL D_6_-glutaric acid (C/D/N Isotopes Inc, D-5227) was added. Cells were scrapped into a pointed microcentrifuge tube kept on ice, and vortexed for 5 s each. Cells were then centrifuged at 20,000 x g for 10 mins at 4 °C, after which 200 µL of the supernatant was removed and transferred to a wide bottom Eppendorf safe-lock tube (Merck; EP0030120094). Samples were then dried in a speedvac system (Thermo Scientific, Savant SPD1010) for 2 h at RT. The dried metabolite pellet was then stored at -80 °C until derivatisation. *Derivatisation*: Pellets were thawed on ice and 50 µl of LC-MS grade water was added, followed by 50 µL each of freshly prepared 3NPH (33 mg/mL in 75% LC-MS-grade methanol and 25% LC-MS-grade water; Sigma-Aldrich, N21804), EDC (20 mg/mL in 100% LC-MS-grade methanol; Sigma 39391), and pyridine (2.5% v/v in 100% methanol; Sigma-Aldrich, 270970. Each tube was vortexed for a minimum of 15 s to resuspend metabolites and was then returned to ice for 1 h protected from light. Samples were then dried in a speedvac for 2 h at RT. Samples were then resuspended in 50 µL of a mix containing 25% LC-MS-grade methanol and 75% LCMS-grade water, with vigorous vortexing, followed by transfer to LC-MS vials. *Acquisition and Analysis*: LC-MS analysis was performed using a Waters Acuity Premier HPLC system coupled to a Waters Xevo TQ-XS. Analyte separation was achieved using a Waters Premier C18 column (1.7 µm, 2.1 x 150 mm) with 0.01% formic acid (LiChropur®) in LC-MS grade water (mobile phase A) and LC-MS grade acetonitrile (Biosolve) with 0.01% formic acid (mobile phase B). The LC flow was set at 0.35 mL/min and the gradient was held at 99% A for 5 min, 94% A at 6 min and held for 10 min, 90% A at 22 min, 85% A at 25 min, 75% A at 30 min, 70% A at 35 min, 0% A at 40 min and held for 3 min, then returning to 99% A at 43.5 min and stabilising for 1.5 min. Data was post-processed using Skyline (MacCoss Lab). Data normalisation (to internal standard and biomass abundance using sulforhodamine B (SRB) analysis) was performed using an in-house R Shiny application. Metabolite enrichment and volcano analysis was then performed using MetaboAnalyst 6.0 software^39^.

### Glutathione measurement by LC-QqQ

*Extraction*: Immediately prior to extraction, a 2.5 mg/mL N-ethylmaleimide (NEM) in LC-MS-grade water stock solution was prepared. For whole cell measurements, cells were washed with NEM wash solution (0.9% (w/v) NaCl in LC-MS-grade water containing 20% NEM stock solution), extracted using 200 µL of NEM extraction solution (40% LC-MS-grade acetonitrile (ACN; Biosolve, BIO-01204102), 40% LC-MS-grade methanol and 20% NEM stock containing 1 µg/mL D_6_-glutaric acid) and each well scrapped into a microcentrifuge tube. Samples were vortexed for 5 s and centrifugation at 20,000 x g for 10 min at 4 °C, after which 150 µL of supernatant was transferred to a LC-MS vial ready for same-day sample injection. For cytoplasmic and mitochondrial fractions, cells were washed with NEM wash solution followed by rapid separation using digitonin. Briefly, 500 µL of mitochondrial buffer (2 mM potassium phosphate monobasic, 120 mM potassium chloride, 3 mM HEPES, 1 mM ethylene glycol BIS tetracetic acid (EGTA), 3 g/L BSA in dH_2_O, pH 7.2; reagents Sigma-Alrich) containing the optimised concentration of digitonin (100 µg/mL; Sigma-Aldrich, D5628) was added to the cells for 2 min at 37 °C. Following incubation, the buffer was rapidly removed and diluted 1:10 with NEM extraction solution, generating the cytosol fraction. The remaining organelle fraction (predominately containing the mitochondria) was quickly washed with ice-cold NEM wash solution, followed by addition of 200 µL NEM extraction solution and scrapping into a microcentrifuge tube. Both sets of tubes were then processed as described above. *Acquisition and analysis*: Chromatographic separation was executed on a Waters Acquity Premier HPLC system integrated with a Waters Xevo TQ-XS mass spectrometer. Samples were resolved using a Waters Premier BEH Amide column (1.7 µm, 2.1 x 100 mm) maintained at 30 °C. The mobile phase consisted of (A) 20 mM ammonium bicarbonate with 0.1% ammonium hydroxide and (B) 1:9 H_2_O/ACN, delivered at a flow rate of 0.5 mL/min. The gradient profile was programmed as follows: 0–0.5 min (5% A), 1.5 min (6% A), 2.0 min (13% A), 4.0 min (18% A), 6.5 min (35% A), 7.0 min (10% A), 7.5–10 min (Return to 5% A and re-equilibrate). Post-acquisition data processing was conducted in Skyline (MacCoss Lab). Data normalisation (to internal standard and biomass abundance using SRB analysis) was performed using an in-house R Shiny application.

### NAD/NADH and NADP/NADPH ratio measurement using LC-QqQ

*Extraction:*Cells were washed with 0.9% (w/v) NaCl in LC-MS grade water and quickly placed onto wet ice, where 200 µL of prechilled extraction solution (40% LC-MS-grade methanol, 40% LC-MS-grade ACN and 20% LC-MS grade water containing 1 µg/mL D_6_-glutaric acid and 0.5% (w/v) formic acid (Biosolve, BIO-069141A8) was added and cells were rapidly scrapped into microcentrifuge tubes. Samples were left on wet ice for 3-5 min, after which the samples were neutralised by adding 15% w/v ammonium bicarbonate (Supelco, 5.33005), before being transferred onto dry ice for 15 min. Samples were then thawed on wet ice before being centrifuged at 20,000 x g for 5 min at 4 °C. A total of 150 µL of the supernatant was then transferred to LC-MS vials for same-day sample injection. *Acquisition and analysis:* samples were acquired using the same instrument parameters and analysis as outlined in the glutathione section above.

### Glucose consumption and lactate production measurements

Following 24 h incubation at the indicated oxygen tensions, cell media was collected and glucose consumption (mmol/L) and lactate production (mmol/L) was analysed using the Stat Profile Prime CCS Analyzer (Nova Biomedical). A cell-free media control was subtracted from readings to account for starting glucose concentration, after which values were normalised to cell biomass and expressed as relative to control cells at 21% O_2_.

### Isotope tracing by Gas Chromatography-Mass Spectrometry (GC-MS)

*Incubation and extraction*: Cells were plated in complete medium onto 6-well plates and left to adhere overnight. The following day, the media was removed, cells washed with 1x PBS, and media was replaced with DMEM Flux media (Sigma-Aldrich, D5030) containing either 10 mM [U-^13^C_6_]-glucose (Cambridge Isotope Laboratories, CLM-1396) or 2 mM [U-^13^C_5_]-glutamine (Cambridge Isotope Laboratories, CLM-1822-H) and 10% dialysed FBS. Media was also supplemented with the appropriate concentration of unlabelled glucose or glutamine, depending on which tracer was being used. Cells were then incubated for 24 h under standard conditions or hypoxia. Following incubation, cells were washed twice with 0.9% (w/v) NaCl, and the plate was placed onto a metal plate over dry ice, where prechilled solutions of 500 µL methanol, 200 µL LC-MS-grade water containing 1 µg/mL D_6_-glutaric acid and 500 µL of chloroform (Sigma-Aldrich, 650498) was added. Samples were then scrapped into microcentrifuge tubes, vortexed and centrifuged at 20,000 x g for 20 min at 4 °C. The upper phase (containing the polar metabolites) was then carefully transferred to a clean safe-lock tube and dried at RT using a speedvac system for 2 h. Dried samples were then stored at -80 °C until day of analysis. *Derivatisation*: On the day of analysis, dried samples were solubilised using 40 µL 2% methoxyamine hydrochloride (Sigma-Aldrich, 226904) dissolved in pyridine and incubated with 500 rpm shaking for 1 h at 60 °C. Following incubation, 60 µL MTBSTFA + 1% TBDMCS (Restek, 35610) was added, and samples were returned to shaking incubation for an additional hour. Samples were then centrifuged at 20,000 x g for 5 min, and the supernatant was transferred to GC-MS vials ready for sample injection. *Acquisition and analysis*: GC–MS analysis was undertaken using an Agilent 8890 GC and 5977A MSD using a 90 m DB-FastFAME column (0.25 um, ID: 0.25 mm). 2-5 μL of sample was injected in pulsed splitless mode with helium carrier gas set at constant pressure (46 psi), sample inlet was 280 °C. Initial oven temperature was 50 °C, then ramping to 160 °C at a rate of 50 °C/min, followed by a ramp to 230°C at a rate of 10°C/min and holding for 30 minutes, and a final ramp to 350°C at a rate of 1 °C/min with a 16 min hold. The transfer line was held at 250 °C. Analytes were ionised via NCI (CI Gas: Methane, Flow: 40%) and source temperature of 280 °C. Compound detection was carried out in scan mode. All data processing was performed in MATLAB (2024a) using the METRAN script. Mass isotopomer distributions (MIDs) were corrected for natural isotope abundance. Each metabolite in all datasets were normalised to the internal standard and cell biomass using SRB analysis.

### Fatty acid measurement by GC-MS

*Extraction*: Adherent cells were cultured in 6-well plates to 70-90% confluence, with an additional well retained for biomass normalisation by SRB assay. All extraction blanks and samples were processed using glassware wherever possible to minimize fatty acid contamination. Cells were washed once at room temperature with 0.9% (w/v) NaCl prepared in LC-MS-grade water and containing 0.01% (w/v) butylated hydroxytoluene (BHT). Ice-cold methanol containing 0.01% (w/v) BHT and 1 µg/mL D27-myristic acid (Cayman Chemical) as the internal standard was then added at 500 µL per well. Cells were immediately scraped into borosilicate glass tubes on ice, and an extraction blank was prepared from an empty well. LC-MS-grade water (500 µL) and dichloromethane (2 mL) were added sequentially to each tube. Samples were vortexed for at least 15 s and centrifuged at 1,000 × g for 10-15 min at 4 °C. The lower organic phase was transferred with a glass Pasteur pipette to a clean glass tube without disturbing the interphase or cellular debris. Comparable organic-phase volumes were recovered across samples and dried under nitrogen to limit lipid oxidation. *Saponification*: For total fatty acid analysis, dried extracts were saponified by adding 200 µL of freshly prepared saponification solution containing 15% (w/v) potassium hydroxide prepared 1:1 with methanol. Samples were vortexed, tightly capped and incubated at 50 °C for 1-2 h protected from light, with a derivatization blank processed in parallel. After cooling and brief centrifugation, 300 µL of 2 M hydrochloric acid in LC-MS-grade water was added to acidify the samples to pH <5. Fatty acids were extracted by adding 500 µL hexane, vortexing and centrifuging for at least 10 min at 4 °C. The upper hexane phase was transferred to a clean glass tube without disturbing the lower phase. The hexane extraction was repeated once, the upper phases were combined, and the solvent was removed under nitrogen at room temperature. *Derivatisation*: For pentafluorobenzyl (PFB) derivatization, 25 µL of 1% (v/v) N,N-diisopropylethylamine in isooctane and 25 µL of 1% (v/v) 2,3,4,5,6-pentafluorobenzyl bromide in isooctane were added to each dried extract. Samples were briefly sonicated, incubated for 30 min at RT protected from light and dried under nitrogen at RT. The PFB-fatty-acid derivatives were reconstituted in 50 µL isooctane, vortexed and transferred to amber GC-MS vials. *Acquisition and analysis*: Data was acquired using the same parameters as outline in the section above (Isotope Tracing).

### Assessment of lipid peroxidation

Lipid peroxidation was assessed using a modified TBARS assay as an index of malondialdehyde (MDA) formation. Briefly, cells were washed with 1x PBS before being lysed in RIPA buffer (50 mM Tris-HCl pH 8.0, 150 mM sodium chloride, 1.0% Igepal CA-630 (NP-40), 0.5% sodium deoxycholate, and 0.1% SDS) with 1x protease inhibitor cocktail (Sigma Aldrich, P8340) and sonicated (40%, 3 x 15 sec pulses) to ensure complete lysis. Lysates were centrifuged at 10,000 x g for 10 min at 4 °C and the resulting supernatant collected for analysis. An aliquot of the supernatant was reserved for protein quantification using the bicinchroninic acid (BCA; Thermo Fisher Scientifc, 23227) assay. To determine MDA concentration, 200 µL of sample or freshly prepared MDA standard (0-100 µM; hydrolysed TMP; Sigma-Aldrich, 108383) was combined with 200 µL thiobarbituric acid (TBA; Sigma-Aldrich, T5500) reagent and incubated in a water bath for 2 h at 95-100 °C, covered from light. Samples were cooled on ice and centrifuged at 10,000 x g for 10 min to remove debris. Subsequently, 200 µL of each reaction mixture was transferred to a black 96-well plate and fluorescence was measured using a PHERAstar FS microplate reader (BMG Labtech) with excitation and emission wavelengths of 510 and 550 nm respectively. Sample MDA concentrations were determined from a standard curve and normalised to protein concentration.

### Cell biomass and viability quantification by Sulforhodamine B assay

At the indicated time points, cells were fixed by adding 20% (v/v) ice-cold trichloroacetic acid (TCA; Sigma-Aldrich, T0699) directly to the media and incubated at 4 °C for a minimum of 30 min. Wells then carefully washed 3 times with distilled water and left to dry, after which they were stained with 0.4% (w/v) SRB (Sigma-Aldrich, 230162) in 1% acetic acid for 10 min at RT. SRB solution was then removed and wells were washed 3 times with 1% acetic acid. Following drying at RT, SRB stain was dissolved using 50 mM Tris/HCL pH 8.8 and absorbance was quantified at 510 nm using a FLUOstar OMEGA microplate reader (BMG Labtech). For cell proliferation and viability analysis, values were normalised to the time zero condition for each cell line at each oxygen concentration.

### 3D growth measurements

Cells were seeded into ultra-low attachment round-bottom plates (Thermo Fisher Scientific, 174925) and allowed to form spheroids (72 h for U87 and HAP1 cells; 96 h for GBM T1 cells). Following spheroid formation (time zero; T0), brightfield phase-contrast images were acquired every 24 h for up to 168 h using an EVOS M5000 imaging system (Invitrogen; 4x objective). Image analysis was performed using a semi-automated approach in FIJI (ImageJ). Images were calibrated according to scale bar and subjected to Gaussian smoothing (σ = 10) to reduce high-frequency image noise. Spheroid boundaries were segmented using the Wand (Tracing) tool with the region-growing tolerance values selected within a predefined range (40-70) according to spheroid size and image contrast. Tolerance values were adjusted to ensure correct inclusion of the visible spheroid body and exclude phase-contrast halo artefacts. Parameters were validated by comparison with manually determined spheroid diameters across representative spheroids of different sizes and circularity. Spheroid area (µm^2^) was recorded for each spheroid at each time point and normalised to the corresponding T0 measurement. Data are presented as the change in spheroid area relative to T0 and representative images depict the same spheroid over the indicated time course.

### Cell treatments

For 2D experiments, cells were treated with 250 µM temozolomide (TMZ; BioServ, BS-6174P) for 72 h at the indicated oxygen tensions. For 3D experiments, U87 and GBM T1 spheroids were treated with 500 µM TMZ while HAP1 cells received 100 µM TMZ, for the indicated durations. Navitoclax (MedChemExpress, HY-10087) and ABT-737 (Stratech, A8193-APE; Supp. Fig. 7f) were administered at the indicated concentrations (1-50 µM) for 24 h. For 3D treatments, U87 and GBM T1 were treated with 25 µM navitoclax for 96h, whereas HAP1 spheroids were treated with 1 µM navitoclax for 96 h or 168 h or with ABT-737 (1 or 10 µM) for up to 240 h (Supp. Figs. 7g, h, j). DMSO served as the vehicle control in all experiments.

#### Immunohistochemistry

HAP1 SLC25A39^+/+^ and SLC25A39^-/-^ cells were seeded on ultra-low attachment round-bottom plates and left to form for 48 h. Spheroids were then treated with DMSO vehicle or 1 µM ABT-737 for 48, 72 and 168 h. At each time point, 96 spheroids per condition were fixed using 10% neutral-buffered formalin overnight at 4 °C and then transferred to 70% ethanol for storage. Spheroids in solution were then transferred to Tissue Wrap (EBB-0106-10A; CellPath, UK) prior to processing to create a FFPE block. All staining was performed on 4 µm FFPE sections. Sections were stained with haematoxylin and eosin (H & E) using a Leica ST5020 autostainer and Carbonic Anhydrase IX (CAIX; NB100-417, Novus Biologicals), Cleaved Caspase-3 (CC3; #9661, Cell Signaling), Ki67 (#12202, Cell Signaling) and N-Cadherin (13116, Cell Signaling) were stained on a Leica Bond Rx autostainer. FFPE sections were loaded onto the autostainer and underwent dewaxing (Leica, AR9222) and epitope retrieval on board using ER2 buffer (Leica, AR9640). All sections were retrieved at 100°C for 20 min, after which sections were rinsed with wash buffer (Leica, AR9590) before peroxidase block for 5 min using an Intense R kit (Leica, DS9263). Sections were rinsed with wash buffer and antibodies were applied at previously optimised dilutions (CAIX, 1:250; CC3, 1:500; Ki67, 1:1000, N-Cadherin, 1/75) for 30 min before washing with buffer then application of Rabbit Envision (K4003, Agilent) for 30 min. Sections were rinsed with wash buffer before visualisation using DAB. Sections were counterstained with haematoxylin before coverslipping the sections with DPX mountant (CellPath, SEA-1300-00A). Images were captured from the stained sections using a Leica Aperio AT2 slide scanner.

#### Combination treatment with irradiation

Cells were plated, left to adhere and either cultured under standard conditions or transferred to hypoxia. The following day cells were treated with TMZ (250 µM for U87; 500 µM for GBM T1) alone or in combination with 10 µM navitoclax. Corresponding time-zero plates were fixed for SRB analysis. After 24 h, cells received 30 Gy irradiation using the Cell Rad+ Benchtop System (Precision X-Ray) and returned to their respective oxygen tensions. Twenty-four hours post-irradiation, cells were re-treated with TMZ alone or in combination with navitoclax for a further 24 h before fixation for viability analysis. Data are presented relative to the irradiation-only control for each cell line at the corresponding oxygen tension.

### 3D viability assay

Viability was assessed using the Viability/Cytotoxicity Assay Kit (Proteintech, PF00007) with the following modifications. Following treatment, media was carefully removed and replaced with serum-and phenol red-free media containing Calcein AM (4 µM for U87 and GBM T1, 2 µM for HAP1) and propidium iodide (PI; 9 µM for U87 and GBM T1, 4.5 µM for HAP1) for 3 h at 37 °C. Solution was then carefully removed and spheroids were washed twice with serum-and phenol red-free media and imaged using the EVOS M5000 imaging system (4x objective). Cell death within spheroids was quantified from PI fluorescence images (Texas Red channel) using FIJI software (ImageJ). PI images were subjected to background subtraction (1000 pixel rolling-ball radius), kept consistent across all images. Spheroid boundaries were defined from the corresponding Calcein AM images by thresholding with FIJI’s default thresholding algorithm to generate binary masks, from which ROIs encompassing the entire spheroid were obtained. ROIs were transferred to the corresponding PI image, and the integrated PI fluorescence intensity and spheroid area were measured. Background fluorescence was determined from a cell-free region of the same image. Corrected PI fluorescence was then calculated using the formula: Corrected PI fluorescence= Integrated density Spheroid Area Background Mean Intensity. To account for variations in spheroid size, corrected fluorescence was normalised to the spheroid area and subsequently normalised to the mean untreated control value for the corresponding cell line within each biological experiment.

### Statistical Analysis

All statistical tests were conducted using GraphPad Prism 10 software (v10.4.0) and R (v4.3.2). Bioinformatic analyses, including single-cell RNA-sequencing analysis, donor-paired pseudobulk differential expression, scFEA-based metabolic flux inference, survival analysis, functional enrichment, cellular-composition deconvolution, and GSVA, were conducted in R using the packages described above. Unless otherwise specified, all tests were two-tailed, and *p* values were adjusted using the Benjamini–Hochberg method where appropriate. *P* < 0.05 or FDR < 0.05 was considered statistically significant.

Error bars, p-values, statistical tests and what *n* represents are indicated in the figure legends.

